# Parallel Implementation of Smith-Waterman Algorithm on FPGA

**DOI:** 10.1101/2021.07.27.454006

**Authors:** Fabio F. de Oliveira, Leonardo A. Dias, Marcelo A. C. Fernandes

**Affiliations:** Laboratory of Machine Learning and Intelligent Instrumentation, nPITI/IMD, Federal University of Rio Grande do Norte, Natal, 59078-970, Brazil; Centre for Cyber Security and Privacy, School of Computer Science, University of Birmingham, Birmingham B15 2TT, UK; Department of Computer and Automation Engineering, Federal University of Rio Grande do Norte, Natal, 59078-970, Brazil

**Author notes:** These authors contributed equally to this work.

## Abstract

In bioinformatics, alignment is an essential technique for finding similarities between biological sequences. Usually, the alignment is performed with the Smith-Waterman (SW) algorithm, a well-known sequence alignment technique of high-level precision based on dynamic programming. However, given the massive data volume in biological databases and their continuous exponential increase, high-speed data processing is necessary. Therefore, this work proposes a parallel hardware design for the SW algorithm with a systolic array structure to accelerate the Forward and Backtracking steps. For this purpose, the architecture calculates and stores the paths in the Forward stage for pre-organizing the alignment, which reduces the complexity of the Backtracking stage. The backtracking starts from the maximum score position in the matrix and generates the optimal SW sequence alignment path. The architecture was validated on Field-Programmable Gate Array (FPGA), and synthesis analyses have shown that the proposed design reaches up to 79.5 Giga Cell Updates per Second (GCPUS).

## 1 Introduction

In Bioinformatic, the analysis can be divided into three parts called primary, secondary, and tertiary analysis [1, 2]. The primary analysis is responsible for generating genomic data information from biological material. In the primary analysis, the sequencing machines create raw genomic data (or raw data). The raw data is composed of several genome reads.

The secondary analysis involves reads alignment and trimming based on quality, and at the end of this step, a whole genomic is created. Finally, tertiary analysis can be characterized as interpreting results and extracting meaningful information from the data. In this last step, many algorithms and techniques can be applied. Also, many applications are created from these analyses. The tertiary analysis covers various applications, from genome characterization to a vaccine or drug treatment creation [2].

A large amount of raw data has been generated in recent years due to the replacement of Sanger sequencing by Next-Generation Sequencing (NGS), also called High-Throughput Sequencing (HTS) [3, 4]. Each sequencing machine can be created about 7 Tera base pairs (bp) per hour (Tbp/h) [5]. The amount of raw data further increased the occurrence of the COVID-19 pandemic. Disease caused by the SARS-CoV-2 virus has been spreading worldwide and has been declared a pandemic by the World Health Organization [6, 7].

After sequencing reads, alignment methods can be performed to map and determine the evolutionary line of the targeted organism, such as its phylogeny. As a result, it is possible to understand the sample’s action mechanics by comparing them with cataloged samples in existing databases [2, 8]. The most used meta-heuristic alignment method for the sequences is the Basic Local Alignment Search Tool (BLAST) due to its fast processing speed and less memory usage than deterministic alignment algorithms [9].

However, different from the meta-heuristics, deterministic alignment methods offer the optimal alignment for a given input sequence instead of an approximate solution. The main deterministic methods are the Needleman-Wunsch (NW) and Smith-Waterman (SW) algorithms for global and local alignment, respectively [10, 11]. Nonetheless, a significant disadvantage of these algorithms is their slow processing speed and high memory usage due to the computational complexity. For example, SARS-CoV-2, commonly vary from 28k to 31k base pairs (bp) in size. Thus, performing thousands of large-size sequence alignments became a real challenge for extracting information on the raw data.

Thus, it is essential that the processing of algorithms associated with the bioinformatics area cover three critical requirements: high processing speed (high-throughput), ultra-low-latency, and low-power [12–14]. Bioinformatics analysis algorithms are critically dependent on the computational infrastructure to cover high-throughput, ultra-low-latency, and low-power requirements. It can be said that there are three generations of infrastructure used, which are: High-Performance Computing (HPC) [15], Graphics Processing Units (GPUs) [16, 17], and Custom Hardware Architectures (CHA) [18–20].

Genomic analysis solutions associated with the first (HPC) and second (GPUs) generation of computational infrastructure use systems based only on software that can be implemented using only CPUs and GPUs. However, these software-only approaches cannot keep up with the growing computational demands of genomic analysis, given the barriers to reducing latency in large volumes using only CPUs and GPUs. In addition, as the number of nodes grows to handle increasing amounts of data, performance is not scaled linearly [15, 21–23]. The third (CHA) generation of infrastructure has been presenting itself as an exciting alternative to satisfy high-throughput, ultra-low-latency, and low-power requirements [24–29].

To overcome the low-speed processing bottleneck and maintain the optimal alignment of deterministic algorithms, parallel hardware implementations for the SW algorithm have been proposed in the literature. The main platforms used are Field Programmable Gate Arrays (FPGAs), Central Processing Units (CPUs), and GPUs. FPGAs are widely known for their flexibility for parallelization and low-power consumption. An FPGA is a matrix of logic blocks that allows designing different circuits, such as processors, logic circuits, and even algorithm development [27]. FPGA platforms can be categorized as third generation computational infrastructure in bioinformatics, as it is a CHA. Also, the logical blocks within the FPGA are independent, allowing operations to be carried out in parallel and only one clock cycle, unlike CPUs that operate sequentially based on instructions and GPUs that require constant access to memories.

Therefore, this work presents a parallel FPGA design with a systolic array structure to accelerate both the Forward and Backtracking stages of the SW algorithm. The main contributions are high-speed data processing implementation and low memory usage. Thus, allowing high scalability. According to [30], the systolic array is a class of parallel computing architecture that describes an array for dense linear algebraic calculations, proposed by [24]. Its hardware implementation usually uses a pipeline structure, where the data is propagated between Processing Elements (PEs). Besides, its main advantage is to reduce the number of memory accesses throughout the data flow. Hence, systolic arrays simplify the architecture and improve the system’s operating frequency. [31].

### 1.1 Related Works

This subsection briefly discusses hardware-based approaches for the SW algorithm that can be found in the literature, such as implementations in supercomputers [32], GPUs [33–36], architectures based on Resistive Content Addressable Memories (ReCAMs) [37] and FPGAs [34, 38–43].

GPUs are well-known for their high degree of parallelism and computing intensity. However, they have a high cost, significant computing latency, and low energy efficiency. The high computing latency is due to the high number of cores and low cache memory to control these cores. In contrast to GPUs, FPGAs are customizable according to the user’s needs, achieving better computing performance and lower latency [14, 44, 45]. However, FPGA hardware development is usually complex and takes a long time.

In recent years, supercomputers have also become widely used for processing massive data. In [32], a hybrid SW-NW algorithm deployed on the Sunway TaihuLight supercomputer (China’s fastest supercomputer with a peak performance of 100 Peta Floating-point Operations per Second (PFLOPS)) is presented. The SW-NW was implemented in Message Passing Interface (MPI) and Athreads (parallel computing modalities) using the SW26010, a heterogeneous multi-core processor, to achieve good scalability and reduce the processing time. According to the authors, the implementation combines both the local and global alignments of the SW and NW algorithms. A runtime analysis for the P50909 protein was carried out with the SW-NW varying from 1 to 64 nodes, where each node corresponds to a multi-thread processor, reaching 9.85 Giga Cell Updates per Second (GCUPS), a speedup of 15.81 × compared to the runtime of a serially performed single node implementation.

Unlike the conventional platforms previously mentioned, the SW algorithm has also been implemented on ResCAMs, as can be seen in [37]. ResCAMs is a storage accelerator system that allows millions of processing units (PUs) to be deployed over multiple silicon arrays. In [37], the implementation was used to compare the homologous chromosomes between humans (GRCh37) and chimpanzees (panTro4), and the only SW step performed was the building of the score matrix. As a result, their proposal achieved 5, 300GCUPS, a 4.8× speedup over the GPU performance. Besides, it also had a 1.7× better energy efficiency compared to an FPGA implementation.

In [38] an FPGA implementation of the BLASTP, a BLAST heuristic aimed at local alignment of protein sequences, is proposed. The architecture was developing using systolic arrays, and it does not need the neighborhood preprocessing step as the K-mers algorithm. They also used the SW gap concept to maximize the final performance of the system. Despite the good performance achieved by the proposed implementation, it did not offer improvements regarding other works.

In [34], a heterogeneous FPGA architecture for sequence alignment is proposed. Unlike most of the works in the literature, their implementation aims to accelerate the entire SW algorithm with the backtracking process. For this purpose, the architecture can process long strings of data based on parallelism and partitioning strategies; and the backtracking process was performed by dividing the equal parts of the similarity matrix, while the search started from the lower right sub-matrix. The tests were performed for 512 Processing Elements (PEs), reaching 76.8GCUPS at 150MHz and 105.9GCUPS (with external memory) for 200MHz. As a result, a speedup of 3.6× to 25.2× was achieved regarding other SW designs implemented on FPGA and GPUs. Besides, it reached a 26% reduction in power consumption compared to the GPU implementation.

Similarly, more FPGA approaches using systolic arrays for the NW and SW with backtracking sequencing techniques have been proposed, such as [41, 46]. In [41], a VHDL SW implementation, using Dynamic Programming (DP) with approximation correspondences for two different strategies, was proposed. It achieved 23.5GCUPS with speedups between 150× to 400× compared to a 2004-era PC. Meanwhile, in [46], the implementation was based on PEs to perform elementary calculations and a diagonally backtracking search, also developed in VHDL. Comparisons were made with the linear and affine strategies, achieving 10.5GCUPS.

In [42], the SW forward and backtracking processes were implemented in an FPGA. The Qnet structure was adopted for communicating with the FPGA, reaching 25.6GCUPS, a speedup of 300× compared to a desktop computer.

Another FPGA alternative for implementing the SW is OpenCL and OpenMP, as shown in [43]. An OpenCL conversion extension is used to synthesize the algorithm and synchronize tasks in multiple Altera FPGAs through the OSWALD software to accelerate the SW on heterogeneous platforms. Tests were performed using a host, a host with an FPGA, a host with a GPU, a host with a Xeon processor, a single FPGA, and two FPGAs. For a single FPGA, 58.5GCUPS was achieved, while two FPGAs reached 114.7GCUPS.

Therefore, it can be noted from the literature that the key points for a high-performance SW implementation on FPGA are the operating frequency and number of PEs, which in turn are associated with the hardware capacity and design critical path. Thus, we present an FPGA implementation for the SW algorithm using systolic arrays, as in [34, 41, 42, 46].

Our approach performs both the Forward and Backtracking stages of the algorithm. Unlike the approaches in the literature, we obtain the alignment path distances during the Forward Stage processing and the maximum score, reducing the complexity of the Backtracking Stage processing. Memories are used to propagate the distances and maximum score, allowing the Backtracking step to follow the path directly. Thus, our architecture achieves good performance (short critical path), reduced memory usage and high scalability, and prevents memory access overlap latency, even implementing the two stages of the SW algorithm.

## 2 Smith-Waterman Algorithm

Smith and Waterman originally proposed the SW algorithm in 1981 to performs local sequence alignment of nucleotides and proteins in the biological field [11]. The sequence alignment of the SW algorithm includes the Forward and Backtracking stages, which are performed by the calculation results of the alignment similarity score. Besides, the alignment is performed based on two input sequences called query sequence, **q**, and dataset sequence **s**. The query sequence can be expressed by

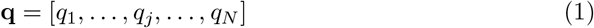

where *q_j_* is the *j*-th nucleotide or amino-acid protein and *N* is the length of the query sequence. The dataset sequence can be expressed by

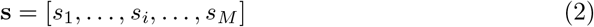

where *s_i_* is the *i*-th nucleotide or amino-acid protein, and *M* is the dataset sequence length. Therefore, the SW algorithm is calculated attractively for two dimensions, and it has a computational complexity of *O*(*M* × *N*).

The Forward Stage calculates the scoring matrix, **H**, where **H** is a two-dimensional array that can only take values greater than or equal to 0 (i.e., 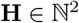). This matrix is generated by comparing the elements of the sequences **q** and **s**. Usually, **H** is generated using DP, and it is initialized with zeroes in the first row and column. Subsequently, the DP process is performed to calculate the sequence scores. Based in works presented in [32, 34, 47], the recurrence relationship can be defined as

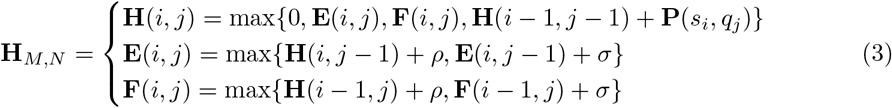

where 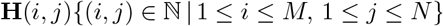, **P** is the score matrix used for obtaining the similarity score between *s_i_* and *q_j_*, **E** and **F** are two assisted matrices when calculating matrix **H**, *ρ* is the gap opening penalty and *σ* is gap extension penalty. In the particular case of *ρ* = *σ*, a linear gap penalty model is obtained, opening and extending a gap with the cost *γ*. **P** is also called a substitution matrix, where the simplest version is when the diagonal receives the match value and the rest of the matrix has a mismatch value. When performing all element calculations, this expression is the **H**_*M,N*_ matrix. Therefore, **H**(*i, j*) is the maximum alignment score of two sub-sequences **s** and **q**. The initialization condition is

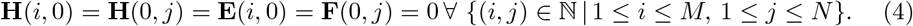

The maximum score value of **H**(*i, j*) in the Forward Stage is the last sequence that will be aligned. To determine the relationship, the previous neighborhood values of the analyzed element are required, i.e., the values on the diagonal, horizontal, and vertical positions, as illustrated in Figure 1. As can be observed, the score of *w* can be found based on its neighborhood (*x,y,v*), which is **H**(*i* – 1, *j* – 1), **H**(*i* – 1, *j*), **H**(*i, j* – 1), respectively. This windowing step occurs throughout the process of determining all scores in **H**.

**Fig 1.**
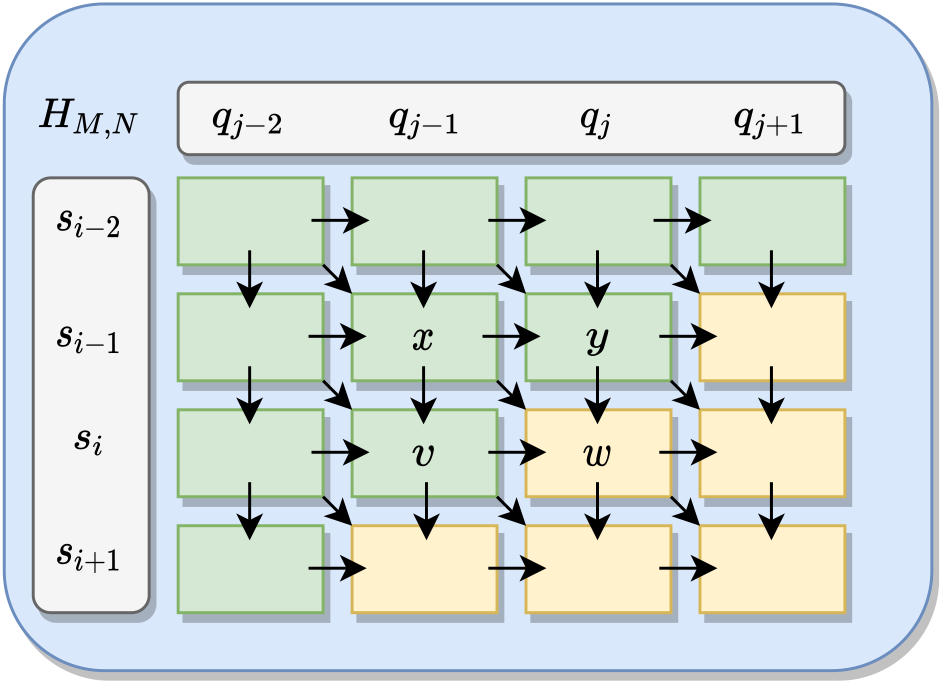
The direction of the score computation in the matrix during the SW Forward Stage. To determine a score, such as w, the neighborhood values (*x, y*, and *v*) have to be known. The green-colored cells indicate already computed values, while the yellow cells indicate that the values to be calculated.

As shown in Figure 1, the neighborhood values *x, y* and *v*, must necessarily be known to determine the value of *w* (i.e., **H**(*i,j*)). For this purpose, those values are defined based on the sequences **q** and **s**. Thereby, the *w* score is determined as

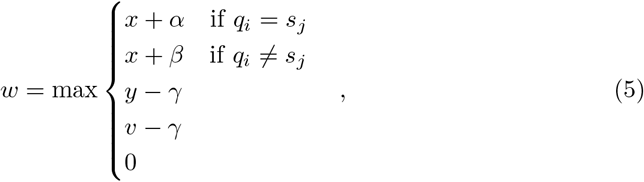

where *γ, α*, and *β* represent the linear gap, a match, and a mismatch, respectively. A gap is a penalty that causes an empty element in the sequence (represented by a dash symbol), while the other sequence continues. It can result from the query or database sequence. The Equation 5 is equivalent the Equation 3, where *x* + (*α* ∨ *β*) = **H**(*i* – 1, *j* – 1) + **P**(*s_i_, q_j_*), *y* + *γ* = **F**(*i, j*) and *v* + *γ* = **E**(*i, j*). Finally, when fully populated, the **H** matrix contains the score and path information.

The Backtracking Stage starts after determining all the scores in the **H** matrix, i.e., calculating the score of all cells **H**(*M, N*). Hence, the backtracking begins at the cell with the highest value in the **H** matrix (maximum score) and trace-back the next position based on the highest neighborhood value, according to Equation 5, which can be on the diagonal, horizontal, or vertical direction. This is an iterative process that repeats until it reaches the limit value, usually set to a score of 0. Also, a directional flag indicates the path. Finally, the backtracking path determines the best local alignment. The diagonal direction points to a match in the alignment, while the horizontal and vertical directions indicate gaps which are represented by dashes in the **s** and **q** sequences, respectively.

## 3 Implementation Description

The hardware architecture for the SW algorithm proposed in this work was developed using systolic arrays to input two DNA sequences and increase the processing speed of the local sequence alignment. An overview of the systolic array structure of the proposal for N PEs is shown in Figure 2. Besides, each PE is divided into 3 modules. These modules are the Forward stage, the storage process, and the Backtracking stage, as seen in Section 2. Each module is illustrated in blue, green, and yellow, respectively. The Forward stage has its module named as Matrix Score Module (MSM), the storage process module is called as Memory Module (MM), and the Backtracking stage has its module as Backtracking Stage (BS).

**Fig 2.**
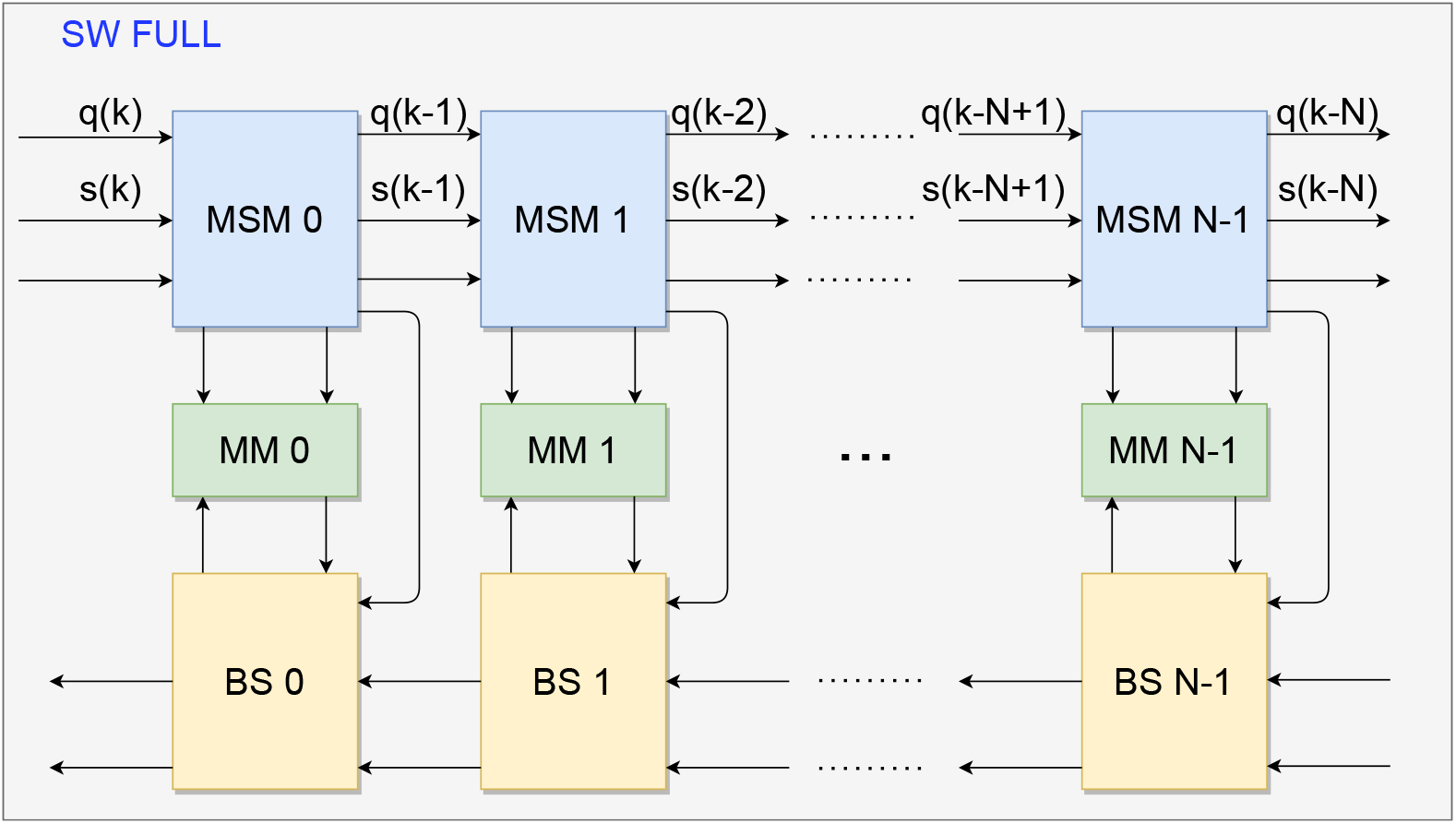
General architecture for the SW algorithm. The Forward Stage (MSM) is represented by the blue block, the Backtracking Stage (BS) by the yellow block, and the Memory (MM) by the green block. Only external signals are displayed, i.e., the **q** and **s** signals.

The labeled signals shown in Figure 2 are generated outside the modules. Meanwhile, the non-labeled ones are generated by computations inside the modules and detailed throughout this Section. The sequences **q** and **s**, defined according to Equations 1 and 2, are external discrete signals used as inputs of the SW algorithm. Furthermore, each signal in the sequences represents one of the four DNA nucleotides, i.e., *A, G, T, or C* (also withstand twenty levels referring to amino acids).

Initially, the circuit starts when the MSM modules propagate the **q** and **s** signals. As seen in Figure 2, each *k*-th element of the **q** and **s** sequences are shifted to each MSM output to shorten and stabilize the critical path, as well as allowing the computation of scores synchronously, preserving the systolic array structure. Afterward, the MSM computes the score according to Equation 3, and propagates the sequence elements to the next MSM; also, the computed results are sent to the respective MM in their order of entry. During this process, the MM operates exclusively in writing mode while the process has not reached the last computation between the two sequences.

The Forward stage is completed after fully computing the scores of the **H** matrix. Also, the last MSM enables the Backtracking process. Consequently, the MM switches to the read mode, and the BS reads the data computed by its respective MSM. The alignment starts from the calculations performed in the MSM. Then, from the respective defined PE in the Forward stage, the process starts and ends according to the definitions of the SW algorithm.

Figure 3 shows the block design that represents each PE of the systolic array, with a detailed description of the signals between the modules within one PE. As can be observed, besides the two input sequences to be compared, **q** and **s**, the MSM also receives an enable signal, **en**. After computing the score between each *k*-th element of the two sequences (i.e., an element of the **H** matrix), the MSM outputs to the next PE the following signals: the calculated score, *Sc_j_*; the maximum score, *MaxVal*, and its position, *AddrRAM*_*i*∧*j*_; the PE *index*; along with the input signals *q, s*, and *en*, shifted in time. In addition, the MSM also outputs signals to the MM, which are the calculated path direction, *Direction*, and the storage address of that path *wAddrDir*.

**Fig 3.**
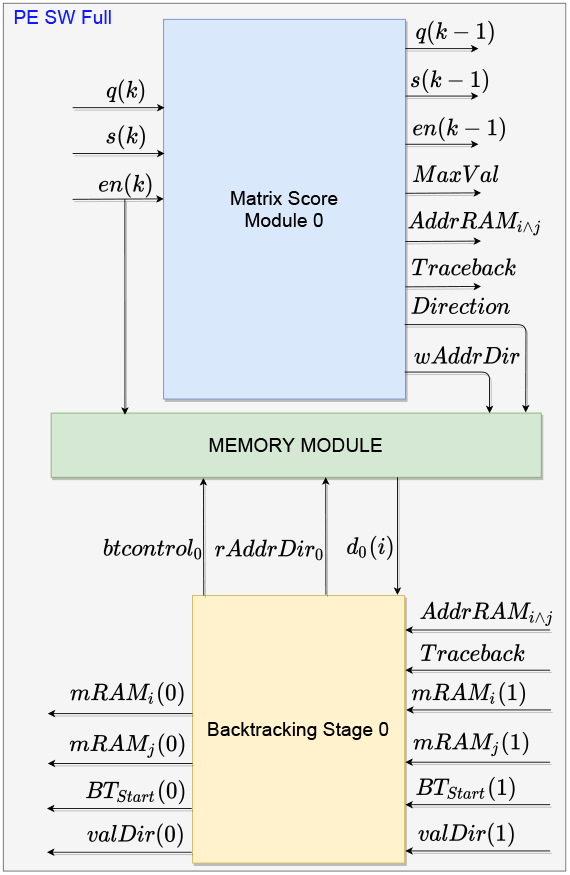
Architecture of each PE in the systolic array. The Forward Stage is represented by the blue block, the Backtracking Stage by the yellow block, and the Memory by the green block.

Subsequently, after fully populating the **H** matrix and, consequently, the **D** matrix, the Forward stage is finished enabling the *Traceback* signal, which in turn begins the BS. Firstly, the BS sets signal *BTstart* to 1, indicating the start of the Backtracking process. Therefore, the *mRAM*_*i*∧*j*_ are propagated back until it reaches the BS with maximum score, which is identified by the signal *index*. From this location match, the *btcontrol* signal is changed to allow the reading of the memory by MM. Thus, the BS receives the path value from the MM at signal *d_j_* when sending the memory address *rAddrDir_j_* signal. The *d_j_* value allows the BS to calculate the next requested address and propagate it to the next module through the *path*(*j*) signal, representing the memory address of the request path in MM. Lastly, the alignment value enters *valDir*, and the process continues until it reaches the complete alignment. All modules are detailed in the following subsections. All signals present in this Section are shown in Table 1.

**Table 1.** Description of signals and the algorithm stage they are used. The Forward Stage is represented by F, Storage Stage by F, and Backtracking Stage by B. They are shown in the Figures 4, 7 and 8.

| Signal | Stage | Description |
| --- | --- | --- |
| $\mathbf{q}(k)$ | F | query sequence, to be compared with the database sequence. |
| $\mathbf{s}(k)$ | F | database sequence. |
| $\mathbf{en}(k)$ | F,S | sequence that enables the PE cells. |
| $\mathbf{Sc}(i)$ | F | vector of scores. |
| $MaxVal$ | F | maximum score. |
| $AddrRAM_{i \wedge k}$ | F,B | memory address of the highest maximum score. |
| $index$ | F,B | corresponding addressing of the modules. |
| $Direction$ | F,S | calculated value of the direction to be stored in RAM. |
| $wAddrDir$ | F,S | storage address corresponding to the direction in RAM. |
| $Traceback$ | F,B | flag to indicate the start of the backtracking in PE $N - 1$. |
| $btcontrol$ | S,B | flag to change the state of the write-to-read memory |
| $rAddrDir$ | S,B | choosing the corresponding value for reading in RAM. |
| $d(i)$ | S,B | return of the value of the path passed from the RAM. |
| $mRAM_{i \wedge k}$ | S,B | addresses of the path to be followed in the alignment. |
| $BT_{Start}$ | B | enable flag of the backtracking after *Traceback*. |
| $BT_{Next}$ | B | flag for enabling the internal circuits to choose and process. |
| $valDir$ | B | alignment path value for that PE. |
| $path(j)$ | B | memory position of the current alignment in the module. |

### 3.1 Forward Approach

Firstly, based on the principles ‘‘divide and conquer” for solving computational problems, we propose a matrix used to store only the values of the recursive path, called the **D** matrix. The **D** matrix is not widely used in the SW literature. However, it is important to achieve a solution at lower-level programming. Besides, a matrix with two different types of information, such as the **H** matrix, increases the hardware design complexity. Matrix **D** needs to store only 4 levels of values which are: 0, 1, 2 and 3. Each element of the matrix **D** needs 2 bits to be expressed, delivering a more economical storage process compared to **H**, which can certainly need more than 2 bits to represent each element.

As previously mentioned, the alignment process is performed based on the query and dataset input sequences, **q** and **s**, respectively. Also, there can have different sizes, represented by N and M, which define the size of the matrices **H** and **D**, respectively. The Matrix Score Module (MSM) calculates the scores and distances in columns of matrices **H** and **D** in parallel.

The systolic array structure developed for the matrices is composed of *N* PEs. Therefore, for each *j*-th element in **q**, there is a *j*-th PE. It is based on dividing the construction of the **H** score matrix expressed by

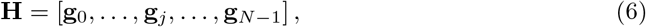

and finding the best path in which the **D** matrix returns the correct sequence alignment, which in turn is equivalent to the directional flags that determined the alignment path. Moreover, for each *PE_j_* (which represents a column of the matrix **H**) there is *i*-th **s**(**i**) that varies from 0 to *M* – 1, according to the following

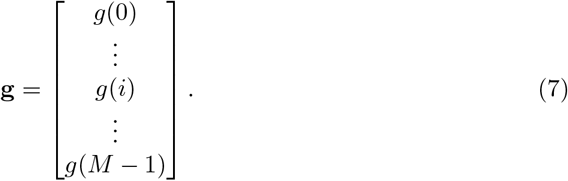

The number of MSMs submodules corresponds to the number of elements in **q**, i.e., 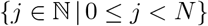, as can be observed in Figure 4. Therefore, **H** is formed by N columns, according to Equation 6. Besides, the MSM also calculates the path, the maximum score value and its position, which are subsequently stored in the Memory Module (MM).

**Fig 4.**
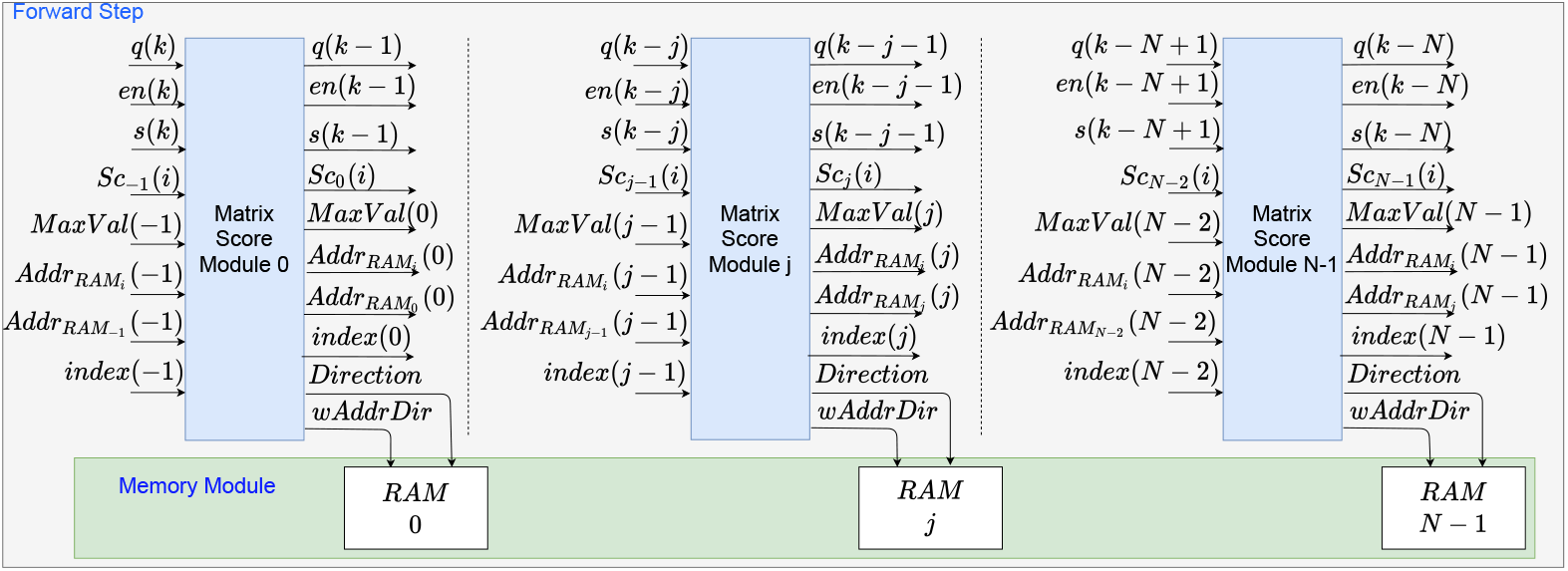
Hardware representation of the **H** score matrix on the Forward Stage. The modules are generated from 0 to *N* – 1.

The SW algorithm in this work is initialized by the *en*(*k*) signal, which enables the memory components in the MSM and MM modules to allocate the two sequences *q*(*k*) and *s*(*k*). The *en*(*k*) is a sequence of pulses of value 1 with size equal to the **s** sequence. Thus, the sequences are transmitted at each sampling time to the Forward Module. The signals are received in MSM, and the respective *q*(*k*) is allocated according to its position, while *s*(*k*) is propagated to the MSM based on the internal counter within each module. The counters within each MSM module are activated with each pulse of the *en*(*k*) signal.

Each *k*-th *q* element is compared to all elements in **s**, iteratively. If the values are equal, a value from the *Match* constant is propagated; otherwise, the value of *Mismatch* is propagated. *Match* corresponds to a reward for similarity, while *Mismatch* is a penalty for inequality between values. Afterward, the addition block sum the values according to

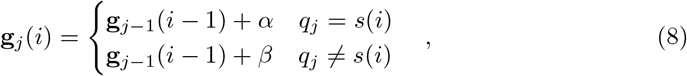

where *α* and *β* are arbitrary values that correspond to the match value and mismatch values, respectively.

Subsequently, the score value, **Sc**_*j*–1_(*i* – 1), and correspondence value, *α* ∧ *β*, are added to define a portion of **g**_*j*_(*i*). The **Sc**_*j*–1_(*i* – 1) value is equivalent to the **H**(*i* – 1)(*j* – 1) value (i.e., **g**_*j*–1_(*i* – 1)). The values of **Sc**_–*j*_, *MaxValue*(–1), *AddrRAM*_*i*∧*j*_ (–1) and *index*(–1) are initialized with 0. At the same time, the **Sc**_*j*–1_(*i*), which is the score value of the previous block, it is received and operated with the value of *Gap*. In addition, the value of the scoring operation of this block in the previous time, *Sc_j_*(*i* – 1), is also operated with the *Gap*. Thus completing the computation of **g**_*j*_(*i*) that can be seen in the Equations 5 and 9.

Figure 5 shows the submodule that constitutes each MSM module. The three blocks in pink are used to perform the addition and subtraction operations, representing the SW’s relations to generate the *M* elements. Thereby, the process of choosing the maximum value among the calculated scores is carried out based on equation 5 as follows

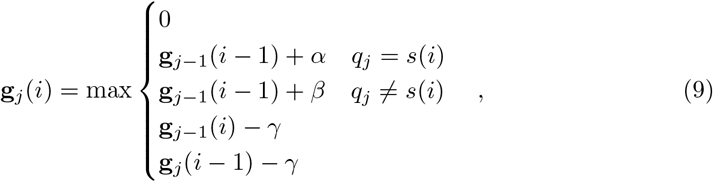

where *γ* is an arbitrary value that represents the chosen linear gap value. This expression is equivalent to Equation 3.

**Fig 5.**
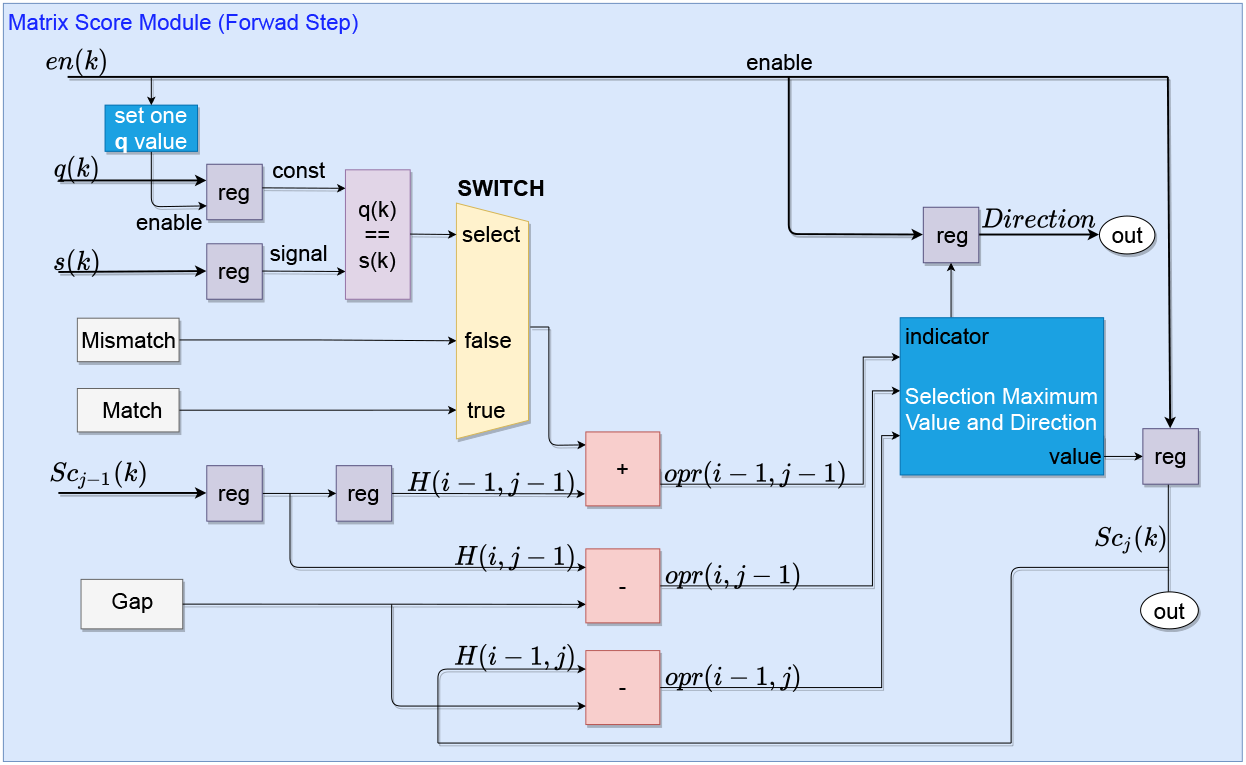
Submodules that constitute a Matrix Score Module. The representation of the circuit and signals is only related to the forward stage.

The output of the pink blocks, called *opr*, are propagated to the next submodule for choosing the maximum score and distance path, as shown in Figure 5. This submodule is built with a set of multiplexers and relational circuits that can find the maximum score value with the coded distance of the path by comparing the *opr* signals, as seen in Figure 6.

**Fig 6.**
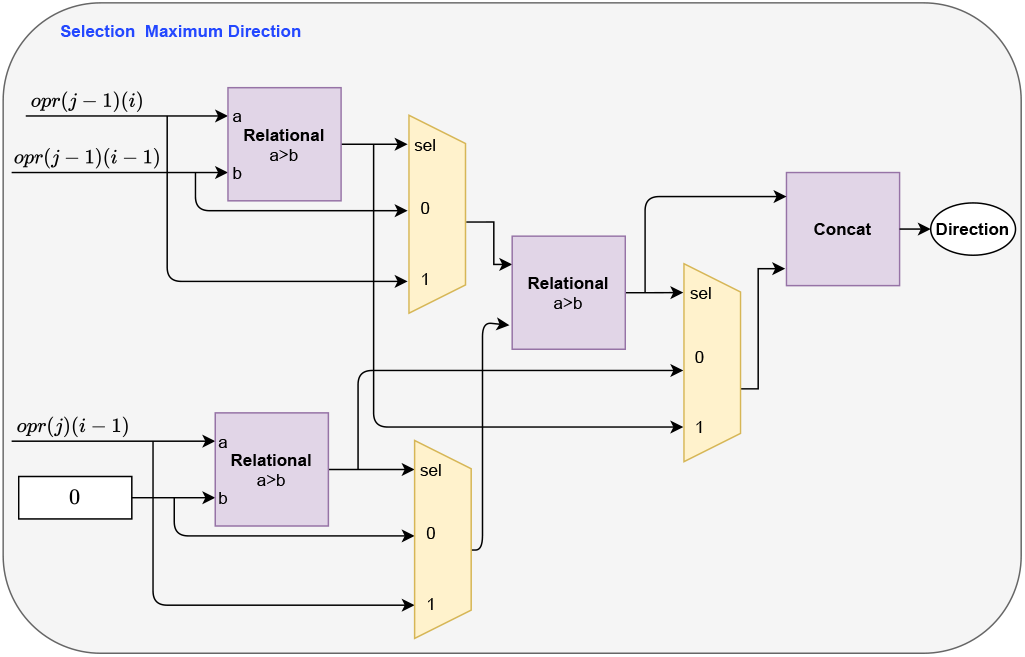
Circuits that constitute the submodule for finding the maximum score and distance path within an MSM. The relational circuits are represented in purple and the multiplexers in yellow.

Selecting path distances is based on a simple encoding of three levels representing the alignment action to be adopted: 2, 1, and 3. Therefore, the levels 2, 1, and 3 represent a match, a gap in the target sequence **q** and **s**, respectively, as described in Section 2. The encoding process of directions is performed in the Forward step, as illustrated in Figure 6. During this process, the same signals used to calculate the **H** score matrix are needed, i.e., the *opr*_*j*–1_(*i* – 1), *opr*_*j*–1_(*i*) and *opr_j_*(*i* – 1), as seen in Figure 5. These values are compared in relational circuits and subsequently chosen according to the criteria of the SW, as seen in the Figure 6.

Then, for demonstrating the realization of the path coding process is done, the information in Figure 1 is used. When looking at the Figure 1, four variables are distributed in an **H** score matrix. The variables *x* = *H*(*i* – 1, *j* – 1), *y* = *H*(*i* – 1, *j*) and *v* = *H*(*i, j* – 1) are known values, while w is a score to be computed. Starting from *w* = *H*(*i, j*) as the observed cell for determining a generic path and *x, y* and *v* as the neighborhood. An integer value is associated with the *d_j_* corresponding to the address of *w*, according to the maximum value determined in the neighborhood, these values are assigned according to the expression

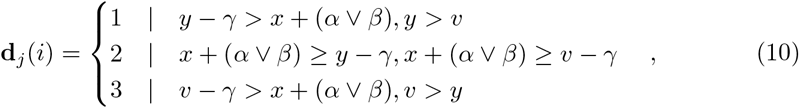

where 1, 2 and 3 is the vertical, diagonal, and horizontal paths, respectively. The Equation 10 is equivalent to the circuit implementation illustrated in the Figure 6, where (*x* + (*α* ∧ *β*)) = *opr*_*j*–1_(*i* – 1), (*y* – *γ*) = *opr_j_*(*i* – 1) and (*v* – *γ*) = *opr*_*j*–1_(*i*). Besides, (*α* ∧ *β*) = *α* for a match and (*α* ∧ *β*) = *β* for a mismatch.

#### Algorithm 1: SW Foward Stage pseudo-code based in structure this proposal

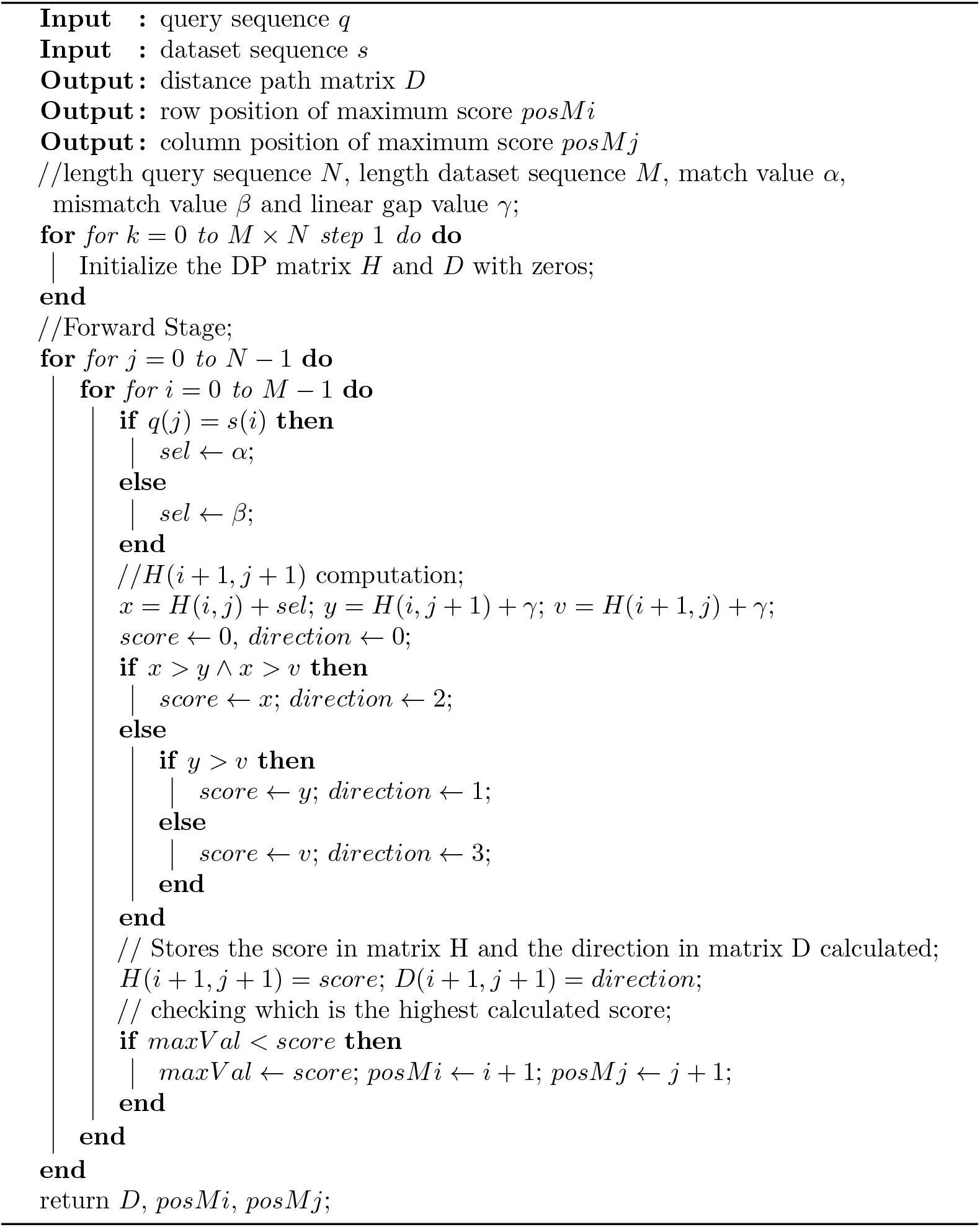

After the process of selection the score and direction, it has the choice of the maximum score based on a logic of multiplexers and relational blocks. There is a counter, called *cntR*, to determines the number of times that the selection of the score and direction is carried out, i.e., the **H** matrix row that the process is on. This is necessary to determine the *AddrRAM_i_* address. At the beginning of MSM processing, *index*(*j* – 1) is added to 1, just once for each MSM, becoming *index*(*j*) and determining the address of this MSM. For the determination of *Maxval*, it is seen whether the previous value is less than the current computed score value, then the calculated current score value becomes the *Maxval, AddrRAM_j_* = *index*(*j*) and respective row process value is *AddrRAM_i_*. It is noted the *AddrRAM*_*i*∧*j*_ signal are corresponding to the location of the maximum score value.

In parallel with the process of determining the maximum score value, there is the process of storing the directions. Thus, the output *Direction* of the submodule is prepared in set with the value *wAddrDir*, which comes from the **H** matrix row calculated at that moment, allowing to write in order in RAM memory according to the respective positions of **H** matrix (i.e., same position of **D** matrix).

Finally, according to the systolic structure, after the MSM processing is over, the signals are parallelly sent to the next MSM. Thereupon, *q*(*k*), *s*(*k*), and *en*(*k*) are shifted in time, that is, *q*(*k* – 1), *s*(*k* – 1), and *en*(*k* – 1), to match the calculation structure of the **H** matrix, as seen in the Figure 4. Besides, the calculated signals *Sc*(*i*), *MaxVal, AddrRAM*_*i*∧*j*_ and *index* are also propagated to the next MSM to preserve the scores calculating structure. This process repeats until the last element of **s** is calculated with the last element of **q**; a counter in is used to determine that moment since the values of the sequences are previously informed to all PEs. The Forward stage finishes with the calculation of the last element of the matrix, i.e., **H**(*M* – 1)(*N* – 1). Consequently, the signal *Traceback* is enabled, indicating the end of the process in all MSM, and the addresses *AddrRAM*_*i*∧*j*_ corresponding to the maximum score value is sent to the next step (i.e., Backtracking process).

Algorithm 1 presents the SW pseudo-code for Forward stage and storage process structures. The Algorithm 1, is prepared to perform the calculation of scores and storage of matrices **H** and **D**. The input is the signals **q** and **s**, which is Equation 1 and 2, respectively. The first loop, in the Algorithm 1, represents each *N* element used, as seen in Figure 2. The second Loop is the interactions made by the signal *En* to allow the calculation of each element of s in each PE. The first conditional structure is the multiplexer for making choices in the MSM, as seen in Figure 5. Submodule Selection Maximum Value and Direction, Figure 6, is represented by the second conditional structure, which compares variables *x*, *y* and *v*. The outputs are **D** matrix stored in MM and the position of the maximum values defined in MSM.

### 3.2 Memory Module (MM)

The MM communicates with both the MSM and the BS, as shown in 7. During the Forward Stage, the data regarding the distance values are written to the MM. Meanwhile, during the BS, the memory addresses to align the sequences are fetched from the MM. The size of each memory is defined by the size of the **s** sequence; also, there is a flag to indicate that the memory is in write mode while computing the **H** matrix and, subsequently, in fetch mode, in the backtracking process.

The MM consists of Random Access Memories (RAMs) used to store the path directions, *Direction*, obtained in the MSM that is thereafter needed in the BS module. Hence, the RAMs are in write mode throughout the Forward Stage and reading mode during Backtracking. The RAM input ports are the address and data busses and write enable mode. Besides, the memory size of each memory is defined based on the size of the sequence **s**, which in turn, the amount of RAM memories is equal to the number of PEs in the systolic array.

**Fig 7.**
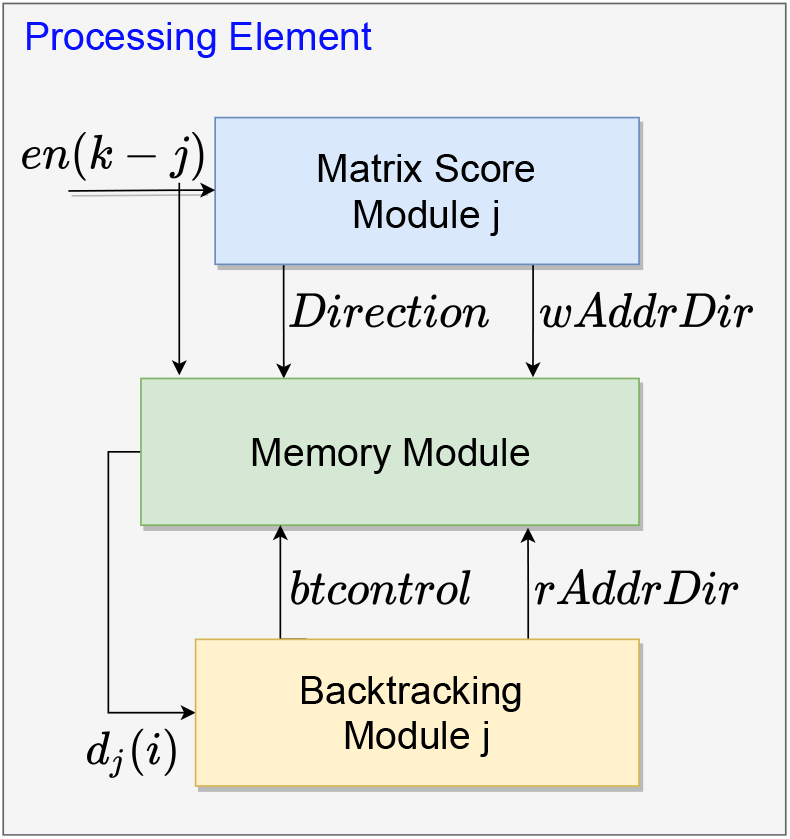
Representation of the simplified Memory Module structure. This model is practically as is the complete processing PE of each column of the **H** matrix.

The enable signal, *en*, is used as write enable for each RAM in the MM. Therefore, *en* =1 defines the write mode, while *en* = 0 the read mode. In addition, the btcontrot signal selects which module controls the RAM address bus. Hence, for *btcontrot* = 0 the memory addresses are defined by the MSM module through *wAddrDir* signal, while *btcontrot* = 1 selects the BS module to define the addresses via *rAddrDir* signal.

Thus, in write mode (*en* = 1 and *btcontrot* = 0) the *wAddrDir* signal defines the address of the RAMs where the *Direction* value is stored by the MSM. Subsequently, after the **H** matrix is fully calculated, the Traceback is enabled to indicate the end of the Forward Stage, and the MM goes into reading mode (*en* = 0 and *btcontrot* = 1). Accordingly, the rAddrDir signal defines the address space the BS fetches the data corresponding to the value reported by the trace-back.

### 3.3 Backtracking Approach

The backtracking process starts when the Traceback signal is enabled in the MSM by counters that determine the last PE and the last processed element of **s**, as described in Forward Stage. As previously mentioned in subsection 3.1, the MSM propagates to the MM the maximum score address that is used as the starting point for alignment, as shown in Figure 8. Meantime, the Figure 9 details the submodules used to create each BS module. The submodules in green are circuits for controlling and synchronizing all signals during the module operation, while the blue submodule performs the alignment path described in this section.

**Fig 8.**
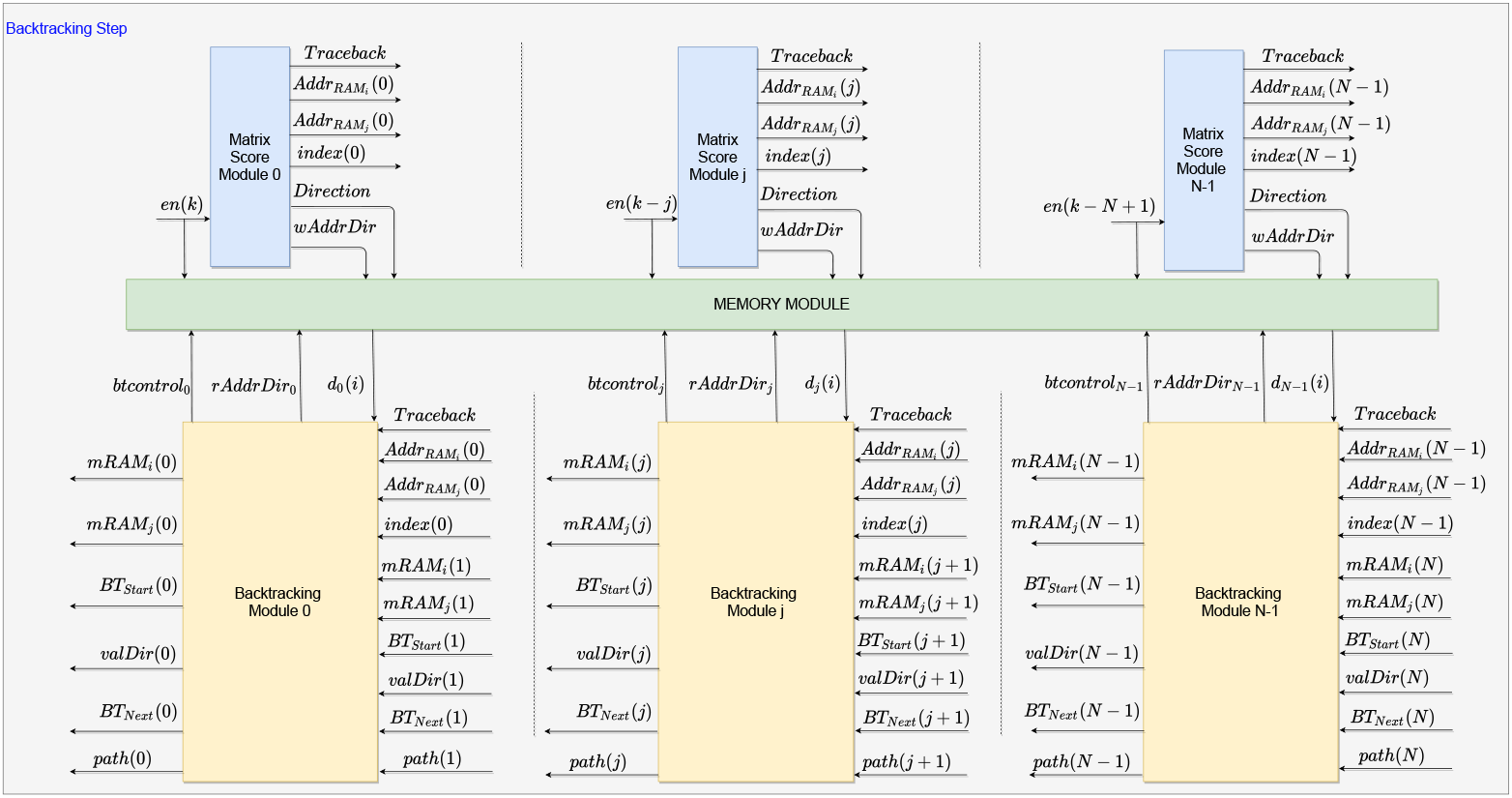
Backtracking Module structure in the FPGA. The operation of this block starts after the Forward Step.

**Fig 9.**
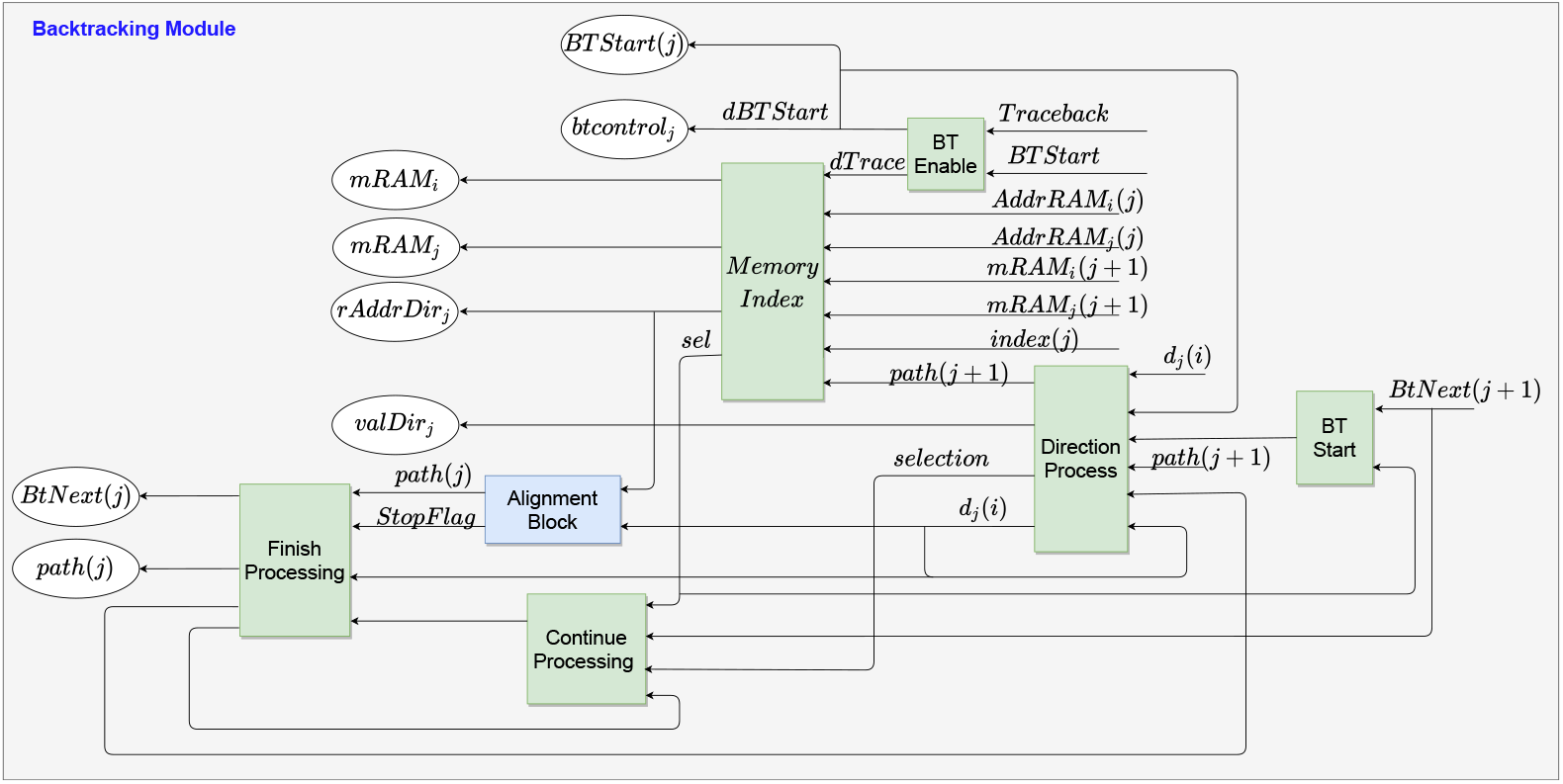
Submodules that constitute the Backtracking Stage Module. The green submodules represent the control submodules, while the blue submodule represents the circuit that performs the alignment.

Firstly, after *Traceback* is enabled, the *BTStart* signal is enabled, and the addresses of the maximum score element, *Addr_RAMi_*(*N* – 1) and *Addr_RAMj_*(*N* – 1), are sent to the respective BS. Also, the values of *Addr_RAMi_*(*N* – 1) and *Addr_RAMji_*(*N* – 1) are assigned to *mRAMi*(*N* – 1) and *mRAMj*(*N* – 1), respectively, by the BT Enable submodule. It is important to emphasize that if the *mRAMj*(*N* – 1) value (i.e., *Addr_RAMj_*(*N* – 1)) is not already in the BS PE, it will trace-back by checking the Memory Index submodule. This process happens until it reaches the PE corresponding to the maximum score location. Afterward, the Memory Index submodule assigns *mRAMi* value to *rAddrDir* to read the memories in the MM, which in turn, returns the *d*(*i*) value to the Direction Process submodule, as can be seen in Figure 9.

Secondly, the alignment process starts. The circuits used to build the alignment submodule are shown in Figure 10. As can be observed, the input *d_j_*(*i*) is used as the multiplexer selector to perform the Equation 10. Therefore, for *d_j_*(*i*) = 3, BS remains in the same memory position and moves back one BS module, i.e., horizontal displacement. While for *d_j_*(*i*) = 1, only the memory position decreases by 1, and BS is verified by the Direction Process and Continue Processing submodules (i.e., vertical displacement). Meanwhile, for *d_j_*(*i*) = 2, the memory position also decreases by 1, and it moves to the previous module with the displacement in the memory position. The circuit after the first multiplexer prevents negative addresses in the memory.

**Fig 10.**
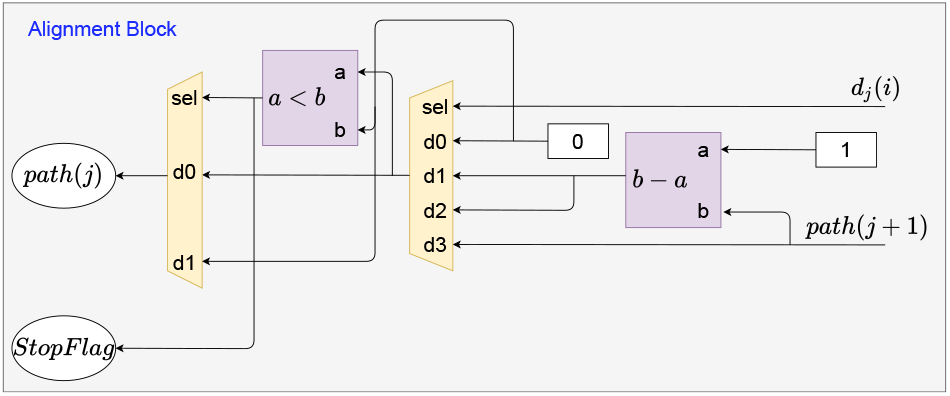
Logical circuits used to build the Alignment Block submodule.

Given that the path to align the first element is found, the Alignment Block submodule receives the *rAddrDir_j_* and *d_j_*(*i*) signals to define the path to be followed by the next BS, as seen in Figure 9. Initially, a logical circuit enables the BT Start and Direction Process submodules to propagate those signals to the Alignment Block. The Direction Process and Continue Processing submodules carry out checks to define which BS module is active, that is, for *d_j_*(*i*) = 1, the data processing is held in the current BS module, and for *d_j_*(*i*) ≠ 1, the signal *BT_Next_* is enabled, indicating the end of data processing in the current PE to start in the next one.

After finding the module for the maximum score, the *mRAM_i_* and *mRAM_j_* signs finish their function. Thus, from the determination of the BS with the maximum score, the *path*(*j*) sign is used as a guide for locating the alignment of each module. Then, the data in MM is requested and the *d_j_* value is returned for verification and establishment of alignment. The verification and establishment of the alignment path is done by the Memory Index, Direction Process, and Continue Processing submodules. Decisions related to *d_j_* value are made in Alignment Block submodule, as illustrated in Figure 10.

Finally, the Finish Processing and Continue Processing submodules finish the data processing in the module. Thereby, the va/Dir output of each submodule is used to construct the alignment path, along with the maximum score position values. The trace-back continues until it reaches *PE*_0_ or finds a path direction with a value of 0.

Algorithm 2 presents the SW pseudo-code for Backtracking stage this proposal. The Backtracking stage, Algorithm 2, is ready to perform the alignment in a list using the path informed in **D**, starting from the positions of the maximum score, as seen in this Section. Inputs for this step are provided by Algorithm 1. The loop for this step represents all Backtracking stage modules from N — 1 to 0. The conditional structure of Algorithm 2 is the representation of submodule Alignment Block, Figure 10, which allows it to trace-back. And the return of the alignment path is storing the data, va/Dir, in RAM memory.

#### Algorithm 2

SW Backracking Stage pseudo-code based in structure this proposal

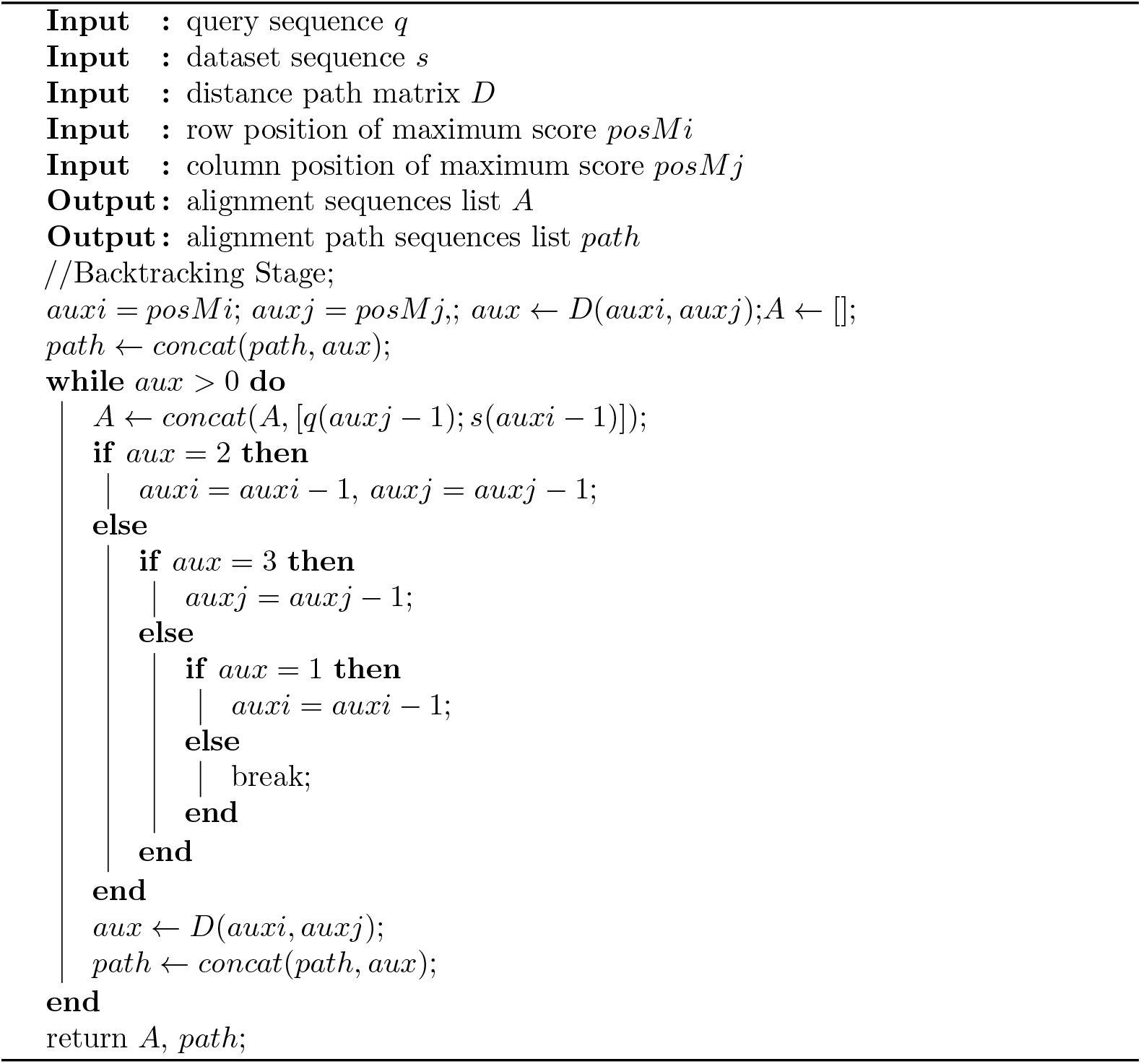

## 4 Results and Discussion

This section presents the synthesis results for the architecture described in the previous section and analyses it regarding the following key points: critical path, operation frequency, number of PEs, and performance. The performance measures the time to calculate an element of the scoring matrix.

The development of the algorithm was carried out using the development platform provided by the FPGA manufacturer, in this case, Xilinx [48]. This platform allows the user to develop circuits using the block diagram strategy instead of VHDL or Verilog. The architecture was deployed on the FPGA Virtex-6 XC6VLX240T and compared to state-of-the-art works. Usually, hardware implementations of the SW algorithm in the literature were implemented only the Forward Stage or both the Forward and Backtracking Stages. In our proposal, both stages were implemented.

The performance for hardware implementations of the SW algorithm is usually measured in Giga Cell Update Per Second (GCPUS), which in turn is defined as

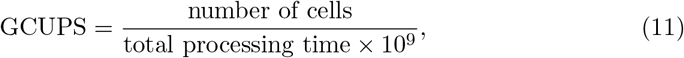

in which a cell can be one vector or matrix element to be computed. This metric can also be described based on the clock frequency, that is,

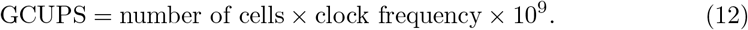

The latter equation is often used to compare the systolic array efficiency. Since the number of cells is equivalent to the number of PEs, and the clock frequency defines the operating frequency, it is unnecessary to measure the total runtime of the algorithm.

### 4.1 Hardware Architecture Validation

To validate the architecture proposed in this work, the sequences **q** and **s** were randomly generated and varying the match, mismatch, and linear gap values. The analysis was carried out for 8 PEs, and **q** and **s** size varied from 8 to 32.

Firstly, the correctness of the matrices **H** and **D** was verified by monitoring the MSM outputs, such as *Sc* and *Direction*, as described in Section 2. Secondly, it was verified if the **D** matrix elements were stored in the correct memory positions in the MM. Lastly, the operation of the BS modules was also verified by monitoring the *path*(*j* + 1) bus and the Memory Index submodule.

Following, the Alignment Block and Direction Process are observed to check if the memory accesses are in accordance with the *path*(*j* + 1) value, that is, according to Equation 10. Also, the Finish Processing and Continue Processing submodules are monitored to verify the values propagated for a match (2), horizontal gap (3), and vertical gap (1).

The data bit-width was defined by the maximum size of the input sequences, limited by FPGA memory capacity. Hence, the input sequence bit-width was set to 3 while constants were defined according to its value. Besides, the bit-width for the MSM buses that perform mathematical operations was defined as log_*total–PEs*_ ×*α*. Meanwhile, the sequence counters for **s** is log_*s–size*_.

Figure 11 shows the architecture deployed and running on the Virtex-6 FPGA. The host computer (i7-3632QM CPU and 8GB of RAM) was used to plot the results and compare them to a software implementation presented in [49], as shown in Figure 12. In the Figure 12, it can be seen that the y axis refers to the **s** sequence, while the x axis refers to the **q** sequence. To increase the resolution of the image, only the parts of the sequences that are aligned are used, where the position at which the alignment starts and the maximum score value are shown in the title of the illustration. The value of Row refers to the position in the **s**, whereas Column is related to the element of the **q**. The amount of sequence alignment performed is represented by Number of Alignments.

**Fig 11.**
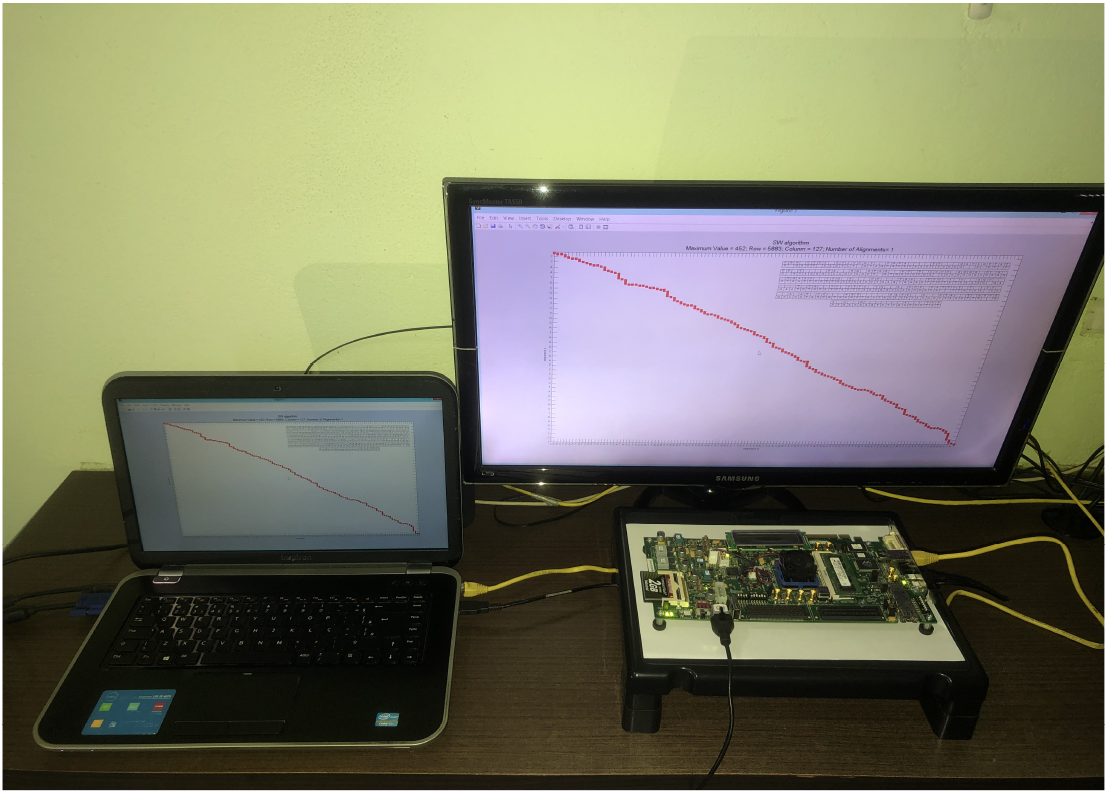
Photo of the hardware architecture deployed on the Virtex-6 FPGA and the host computer used to plot the results.

**Fig 12.**
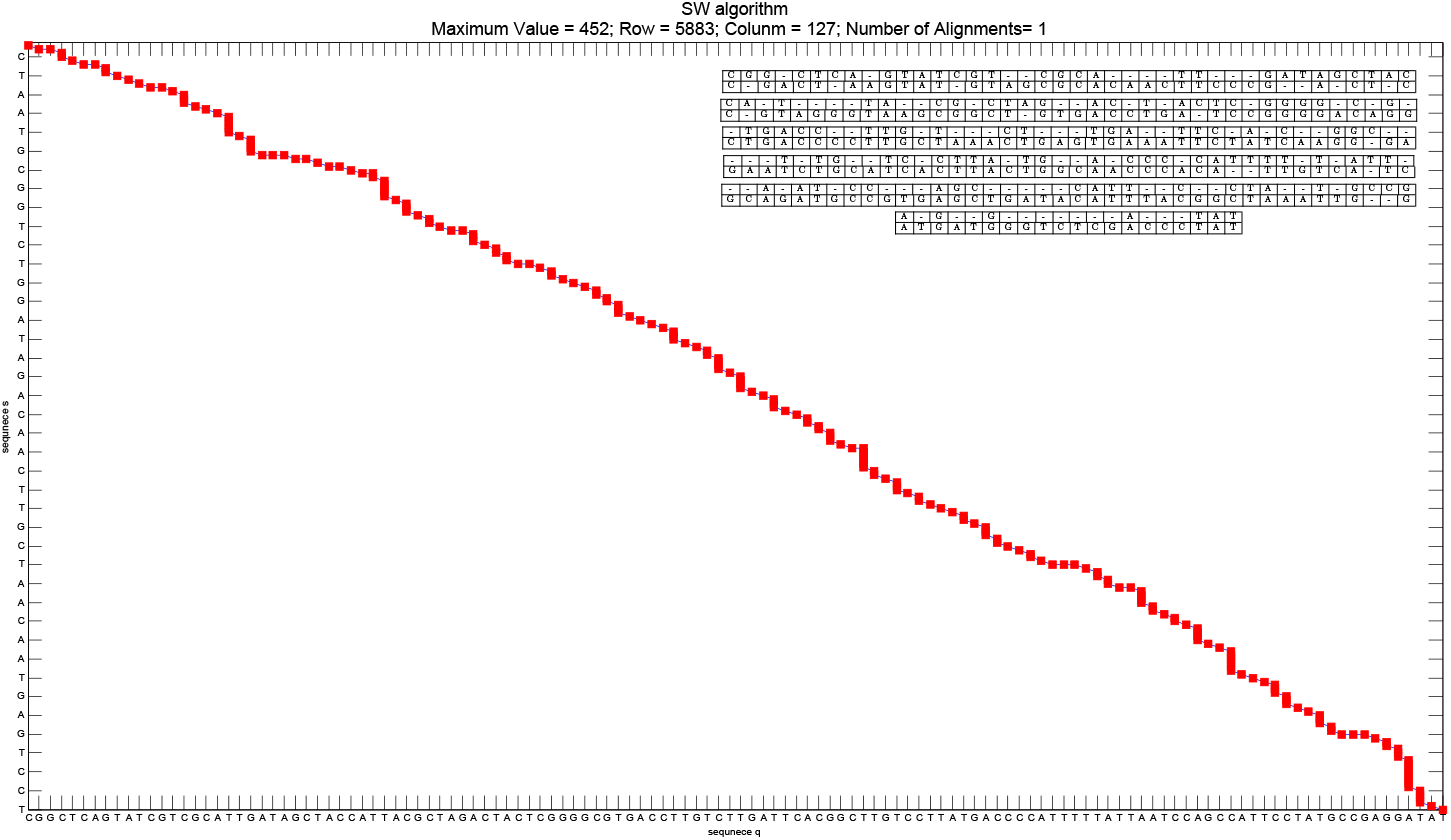
Illustration of the results obtained from our proposal in co-simulation. The image is the most detailed representation of the monitor in Figure 11. It can see that the y-axis refers to the **s**, while the x-axis refers to the **q**. The position at which the alignment starts is indicated by Row and Column. The maximum score value found is presented by Maximum Value. The amount of sequence alignment performed is represented by Number of Alignments.

The architecture parameters for the demo were set to *match* = 5, *mismatch* = –5, *gap* = 1, and 128 PEs. Hence, the size of the sequence **q** is also 128. Meanwhile, the size of the sequence **s** was set to 8,192, resulting in a total of 1,048, 576 calculated cells. Sequence **q** is loaded into memory at each iteration, where it can vary between 4 different 128 nucleotide sequences in the demonstration. The demo is available at [50].

### 4.2 Synthesis Analysis

Analysis of the synthesis results for the SW hardware implementation were carried out for two FPGAs: Virtex-6 XC6VLX240T and Virtex-7 XC7VX485T. Table 2 presents the hardware area occupation and frequency for a different number of PEs. The size of the input sequences were defined according to the number of PEs.

**Table 2.** Area occupation results based on the FPGA synthesis of our SW implementation, with forward and backtracking stages.

| FPGA Model | Array Size (PE) | Slice LUTS / (%) | Memory / (%) | Frequency (MHz) |
| --- | --- | --- | --- | --- |
| Virtex-7 | 512 | 103,778 (34%) | 8,192(6%) | 155 |
| Virtex-6 | 512 | 103,807 (68%) | 8,192(14%) | 120 |
| Virtex-6 | 256 | 47,725 (31%) | 2,048 (3.5%) | 112 |
| Virtex-6 | 128 | 24,386 (16%) | 512 (1%) | 117 |
| Virtex-6 | 64 | 10,803 (7%) | 128 (0.2%) | 157 |

The critical path of the design was ≈ 8.34ns and ≈ 6.44ns for the Virtex-6 and Virtex-7, respectively. Therefore, the maximum clock frequency was 120MHz for the Virtex-6 and 155MHz for the Virtex-7. Regarding the FPGA area occupation, increasing the number of PEs also increases the hardware resources used. For 512 PEs in the Virtex-6, a total of 68% of the Slice Look-Up Tables (LUTs) were used in contrast to only 7% for 64 PEs. Concerning the frequency, a slight decrease is observed as the number of PE increases due to an increase in the critical path. Concerning the Virtex-7, there are unused FPGA resources as less than 35% of Slices LUTs were used. Therefore, it can be used to increase the number of PEs and, thus, the performance.

### 4.3 Comparison with other works

Comparisons with state-of-the-art works were also performed. The performance of systolic array-based implementations increases with the number of PEs. Hence, the comparisons were carried out for the maximum number of PE in each proposal.

The works presented in Table 3 are available in [34]. The second column indicates whether the backtracking stage was also developed on FPGA or only the forward. Meantime, the third to fifth columns present the number of PEs, operating frequency, and performance, respectively. The performance was obtained according to Equation 12. As can be seen, our approach and the one proposed by [34] were the only ones to implement a high number of PEs. However, in [34] only the backtracking path was deployed on the FPGA, and a submatrix structure is used to load the path chosen for alignment. Meanwhile, our architecture relies on a memory storage structure and the definition of the maximum score to align the sequences.

**Table 3.** Table adapted from paper [34]. it compares the proposed SW using reconfigurable hardware based on the operating frequency, number of PEs and performance in GCUPS. It also shows if the works use backtracking or not in the implementation. * indicates the approach uses external memory to accelerate the alignment process.

| Related Works | Backtracking (Yes or No) | PE number in array | Frequency (MHz) | Performance (GCUPS) |
| --- | --- | --- | --- | --- |
| 2005 [51] | No | 252 | 50 | 13.9 |
| 2006 [41] | Yes | 303 | 77.5 | 23.5 |
| 2007 [52] | No | 384 | 66.7 | 25.6 |
| 2007 [53] | No | 128 | 125 | 16.0 |
| 2008 [42] | Yes | 256 | 100 | 25.6 |
| 2009 [46] | Yes | 168 | 62.5 | 10.5 |
| 2011 [54] | No | 100 | 111 | $11.1 \times 12$ |
| 2012 [55] | No | – | 250 | – |
| 2012 [56] | No | $< 100$ | 125 | $16 \times 8$ |
| 2012 [57] | No | 100 | 175 | 17.5 |
| 2012 [47] | No | 128 | 60 | 7.62 |
| 2014 [58] | No | 200 | 200 | 40 |
| 2017 [34] | Yes | 512 | 150 | 76.8 |
| 2017* [34] | Yes | 512 | 200 | 105.9 |
| This work | Yes | 512 | 155 | 79.5 |

Furthermore, a comparison with [34] was also carried out regarding the FPGA area occupation, and it is presented in Table 4. The second and third columns present the FPGA and the number of PEs used, respectively. Meanwhile, the third and fourth columns present the slices and memory blocks occupied, and the fifth column the operating frequency.

**Table 4.** Table with the summaries of the results of the FPGA synthesis works of SW implementation (hardware SW with backtracking step). The Slice column is related to the logical distribution and refers to the occupied slices in the synthesis.

| Related Works | FPGA Model | Array Size (PE) | Slices / (%) | Memory / (%) | Frequency (MHz) |
| --- | --- | --- | --- | --- | --- |
| This work | XC7VX485T | 512 | 35,286/(46%) | 0 BRAM/(0%) | 155 |
| 2017 [34] | XC7VX485T | 512 | 57,870/(76%) | 896 BRAM/(87%) | 200 |

As shown in Table 4, for the same number of PEs, our architecture occupied 35, 286 slices and 0 BRAMs in contrast to 57, 870 slices and 896 block RAMs (28 Mbits memory) in [34]. Also, the total area occupation was higher than 60%, compared to 46% on ours, due to the substitution matrix. Therefore, our proposal has high scalability due to the low resource usage (can reach up to 1, 024 PEs for the XC7VX485T). Besides, our implementation proposal can be implemented in smaller FPGAs, such as the Virtex-6 XC6VLX240T, with a reasonable nucleotide sequence.

Regarding the operation frequency, our proposal can reach up to 155 MHz. So, it is observed that the proposals with the best performances have a similar structure, even with different approaches to the solution. Our proposal and [34] achieving the same performance for the frequency of 150 MHz.

Therefore, our work uses fewer hardware resources to perform the alignment process due to the chosen backtracking approach. As the backtracking stage results in high computational complexity, we simplified the process using the path mapping through the maximum value in **D** and **H**, resulting in linear computational complexity. On the other hand, the architecture proposed by [34] uses considerably more memory resources due to data partitioning and prefetching for the backtracking step. Despite both works achieving similar performance due to the systolic array, there are significant differences in the alignment approach chosen for the FPGA implementation.

The hardware implementation of the alignment process through our approach, developed based on a chain of directions and the maximum score address, is a key contribution for the low use of memories and, thus, achieve high hardware scalability. Hence, the proposed method can compress the data, using only 3 bits in a fixed-point implementation.

## 5 Conclusion

This paper presented a parallel FPGA platform design to accelerate both the Forward and Backtracking stages of the SW algorithm. The main contributions were the high-speed data processing implementation and low memory usage that allowed high scalability. In order to satisfy the high-throughput, ultra-low-latency and low-power requirements and to alleviate the raw data processing problem in bioinformatics. From the strategy of storing alignment path distances and maximum score position during Forward Stage processing. It was possible to reduce the complexity of Backtracking Stage processing which allowed to follow the path directly. The proposal architecture achieved a satisfactory critical path, reduced memory usage and high scalability for two-step SW algorithm. Synthesis results showed that the proposed method could support up to 1, 024 PEs in only one FPGA, using the Xilinx Virtex-7 XC7VX485T. The main advantage is the low hardware resource usage and high performance of 79.5 GCUPS, with an operating frequency of up to 155MHz, without using external resources.

## Acknowledgments

The authors wish to acknowledge the financial support of the Coordenação de Aperfeiçoamento de Pessoal de Nível Superior (CAPES) for their financial support.

## Notes

### Competing Interest Statement

The authors have declared that no competing interests exist.

